# Estimates of persistent inward currents in human spinal motor neurons: non-linear and sex-dependent trajectories from young to very old age

**DOI:** 10.64898/2026.09.24.754101

**Authors:** C. Chatain, R. Hamard, T. Cattagni, R. Roty, R. Lepers, V. Rozand

## Abstract

The contribution of persistent inward currents (PICs) to spinal motor neurons discharge rate is lower in old compared to young adults. However, whether this difference further progresses at very old age (≥80 years) and differs between males and females remains unknown. Thirty young (26 ± 4 years, 15 females), thirty old (72 ± 5 years, 15 females) and twenty-nine very old adults (85 ± 3 years, 15 females) performed isometric dorsiflexion ramp contractions at 20% of maximal voluntary torque. Individual motor unit spike trains were identified from decomposed high-density surface electromyograms of the tibialis anterior. The paired motor unit technique was used to estimate the prolongation effect of PICs (ΔF and normalized ΔF), and the quasi-geometric approach to estimate neuromodulatory (brace height) and inhibitory pattern (attenuation slope) contributions. ΔF was lower in old and very old adults than in young adults, with no difference between old and very old adults regardless of sex. Compared with young adults, normalized ΔF was lower in both old and very old males, whereas in females it is only lower in the very old group. Brace height and attenuation slope did not differ across ages or sexes. These findings indicate that age-related changes in spinal motor neuron excitability are established by the seventh decade and do not progress further in healthy active very old adults, following a nonlinear trajectory that may differ between males and females. The absence of age-related differences in the neuromodulatory index does not support a lower monoaminergic drive in older adults.

## **I)** INTRODUCTION

Aging is a ubiquitous biological process that results in progressive impairments of the neuromuscular system, contributing to reduced functional capacity, loss of autonomy, and frailty. These impairments do not evolve linearly throughout the lifespan, as illustrated by the accelerated decline in maximal force production capacity observed in very old adults (≥ 80 years) compared to old adults (≥ 60 years) (Skelton et al. 1994; Dodds et al. 2016; Haynes et al. 2020; Varesco et al. 2022). Alongside the decline in muscle mass, neural factors also contribute to the age-related impairments in neuromuscular function (Hunter et al. 2016). In particular, aging is characterized by a progressive loss of motor units (MU) and impaired rate coding (McNeil et al. 2005; Orssatto et al. 2022a). Recent evidence suggests that the latter may partly be explained by changes in the intrinsic excitability of spinal motor neurons (for review, see Orssatto et al. 2023).

Spinal motor neurons constitute the final common pathway for force production (Sherrington 1906). They integrate ionotropic excitatory and inhibitory inputs from several peripheral and central sources through nonlinear processes governed by intrinsic membrane properties, particularly the activation of persistent inward currents (PICs) (Heckman et al. 2005). These depolarizing currents, which are generated by voltage-sensitive channels such as L-type calcium and sodium channels, amplify and prolong the output of motor neurons for a given ionotropic synaptic input (Heckman et al. 2005; Binder et al. 2020). PICs are modulated by monoaminergic drive, especially serotoninergic and noradrenergic projections from the brainstem, and could, under certain conditions at least, amplify synaptic input by as much as five-fold (Lee and Heckman 1996; Binder et al. 2020).

Growing evidence indicates that aging impairs PICs. Studies using the well-established paired MU technique (Gorassini et al. 1998, 2002) have consistently reported lower ΔF values (commonly interpreted as an index of the prolongation effect of PICs) in both upper and lower limb muscles of older adults (Hassan et al. 2021; Orssatto et al. 2021, 2022b; Guo et al. 2024). This age-related difference in ΔF amplitude suggests a diminished capacity of spinal motor neurons to self-sustained firing. Although ΔF primarily quantifies this prolongation effect, it is also commonly used as a global index of PICs contribution to MU discharge behavior. Lower ΔF values have therefore been assumed to reflect an overall weakening of PICs effects, including a reduced amplification of synaptic input that may contribute to the lower MU peak discharge rates observed with aging. This impairment has been proposed to result notably from a decline in monoaminergic drive (Orssatto et al. 2023). However, aforementioned studies present several limitations that should be considered.

First, the existing literature has primarily focused on comparisons between young and old adults with limited representation and no differentiation for very old adults. Such an approach overlooks the non-linear trajectory of neuromuscular aging, as illustrated by the greater loss of MUs at advanced ages, especially after the seventh decade (McNeil et al. 2005; Hepple and Rice 2015). Therefore, characterizing the trajectory of age-related changes in the intrinsic excitability of spinal motor neurons is particularly important in this population nearing functional limitations.

Second, previous studies have been conducted predominantly in males due to technical constraints and the difficulty in detecting sufficient MUs in females (Taylor et al. 2022). However, recent evidences suggest that PICs amplitude, estimated by ΔF, may be modulated by biological sex, with greater contribution of PICs in females compared to males (Jenz et al. 2023; Yacyshyn et al. 2025). This underrepresentation limits the generalization of the age-related difference in PICs to females and precludes a comprehensive understanding of potential sex-related difference in PICs regulation across aging.

Third, most of these studies have relied on the ΔF metric as a global estimate of the contribution of PICs to MU discharge behavior. However, while PICs amplitude is largely determined by neuromodulatory drive, the extent to which PICs are expressed in motor neuron discharge also depends on inhibitory inputs (Orssatto et al. 2022b; Gomes et al. 2024). As ΔF reflects this net expression, it does not allow a dissociation between neuromodulatory and inhibitory contributions (Beauchamp et al. 2023; Chardon et al. 2024). Considering the age-related alterations in inhibitory circuits (Butchart et al. 1993; Kido et al. 2004), it is unclear whether changes in ΔF observed in older adults reflect modifications in monoaminergic drive or arise from alterations in inhibitory patterns or a combination of both. A recent framework developed by Beauchamp et al. (2023) now allows to complement ΔF with a quasi-geometric approach based on nonlinearities of MU discharge patterns during the ascending phase of ramp contractions. Computational models have shown that the neuromodulatory and inhibitory contributions to motor neurons discharge behavior could be dissociated using brace height, which is predominantly sensitive to neuromodulatory input, and the attenuation slope, which reflects the level of synaptic inhibition relative to excitation (i.e., push-pull pattern) (Beauchamp et al. 2023; Chardon et al. 2024). This framework has recently been applied to aging by Connelly et al. (2026), who reported that the lower ΔF observed in the tibialis anterior of older adults was accompanied by preserved brace height but lower attenuation slope, at least at low and moderate contraction intensities. These results thus suggest an altered balance between excitation and inhibition in older compared to young adults, rather than a reduced monoaminergic drive. Whether this pattern persists at more advanced ages and differs between males and females remains unknown.

The primary aim of the present study was to investigate age- and sex-related differences in estimates of PICs and their underlying neuromodulatory and inhibitory contributions across young, old (≥ 60 years) and very old adults (≥ 80 years). We sought to explore (i) whether PICs are lower in very old adults compared to old and young adults, and (ii) whether sex modulates PICs within and across age groups. To address these questions, we examined discharge patterns of a large sample of MUs identified non-invasively through decomposition of high-density electromyographic signals. We hypothesized that PICs would be progressively lower with advancing age, with the lowest values observed in very old adults. Considering evidence that young females exhibit greater PICs estimates than males, and that age-related neuromuscular declines may occur at much greater rates in females with advanced age (Hunter 2025), we hypothesized that sex-related differences in PICs would be reduced with aging.

## **II)** METHODS

### 2.1) Participants

Thirty young adults (15 females; 25.9 ± 4.1 years), thirty old adults (15 females; 71.6 ± 5.1 years) and twenty-nine very old adults (15 females; 84.6 ± 3.4 years) participated in this study. An *a priori* power analysis conducted using G*Power (version 3.1.9.7; Faul et al. 2007) based on a two-way ANOVA design (group × sex) indicated that 83 participants were required to detect a large effect size (f = 0.4) with a probability level of 0.05 and a statistical power level of 0.9. Considering an expected data loss of ∼ 10 % due to potential difficulties in identifying MUs from high-density surface electromyography recordings (Lapole et al. 2023), we planned to include a total of 90 participants in this study (i.e., 30 participants in each group with equal numbers of men and women). Due to the difficulty of recruiting very old adults that are not taking medications directly affecting the monoaminergic system, we stopped the inclusions at 29 participants for this group. Participants were free of lower limb musculoskeletal injuries, had no history of neurological disorders and were not taking medications that could influence the monoaminergic system, such as beta-blockers or serotonin reuptake inhibitors. They were asked to avoid strenuous exercise for 24 h before the experimental session and keep their caffeine consumption habits consistent for the duration of their participation in the study. Old and very old participants were additionally required to have a Mini-Mental State Examination (MMSE) score ≥ 24 to ensure normal cognitive function (Folstein et al. 1975). This study received ethical approval from a local ethic committee (IRB00012476-2025-27-02-381) and was conducted in accordance with the Declaration of Helsinki at the exception of the study registration. All participants provided written informed consent prior to their participation in the study and were free to withdraw at any time.

### 2.2) Study design

Participants visited the laboratory on two separate occasions over a 1-week period. During the first visit, participants were familiarized with the study procedures and equipment, and performed several contractions of dorsiflexor muscles including maximal voluntary contractions (MVCs) and triangular ramp contractions. All participants received an accelerometer to assess their physical activity level. Old and very old participants also performed functional assessments (Timed up and go test and 6-min walk test) and completed the MMSE questionnaire. These measures were collected to provide a comprehensive characterization of participants’ physical and cognitive status.

During the second visit, participants completed a standardized warm-up consisting of three contractions at each of the following intensities: 30%, 50% and 70% of the MVC assessed during the first visit. After a 2-min recovery, participants performed at least three maximal voluntary dorsiflexion interspaced by 1 min of rest to assess maximal voluntary torque (MVT). If the difference between the two best attempts was > 5%, additional MVCs were performed. Strong verbal encouragements were given during each MVC to ensure maximal effort. The highest value obtained was used as a reference for the triangular-shaped ramp contractions. Then, participants performed triangular-shaped ramp contractions to 20% MVT of dorsiflexors while high-density surface electromyography (HDsEMG) was recorded from the *tibialis anterior* muscle of the right leg. These ramp contractions consisted of a 10-s ramp-up phase followed by a 10-s ramp-down phase (torque rise and decline of 2 %.s^-1^) (Figure 1A). This modality of ramp contraction has been used extensively to determine ΔF (Hassan et al. 2021; Orssatto et al. 2021; Goreau et al. 2024; Guo et al. 2024). If the produced torque deviated importantly from the expected triangular shape, additional ramp contractions were performed.

**Figure 1.**
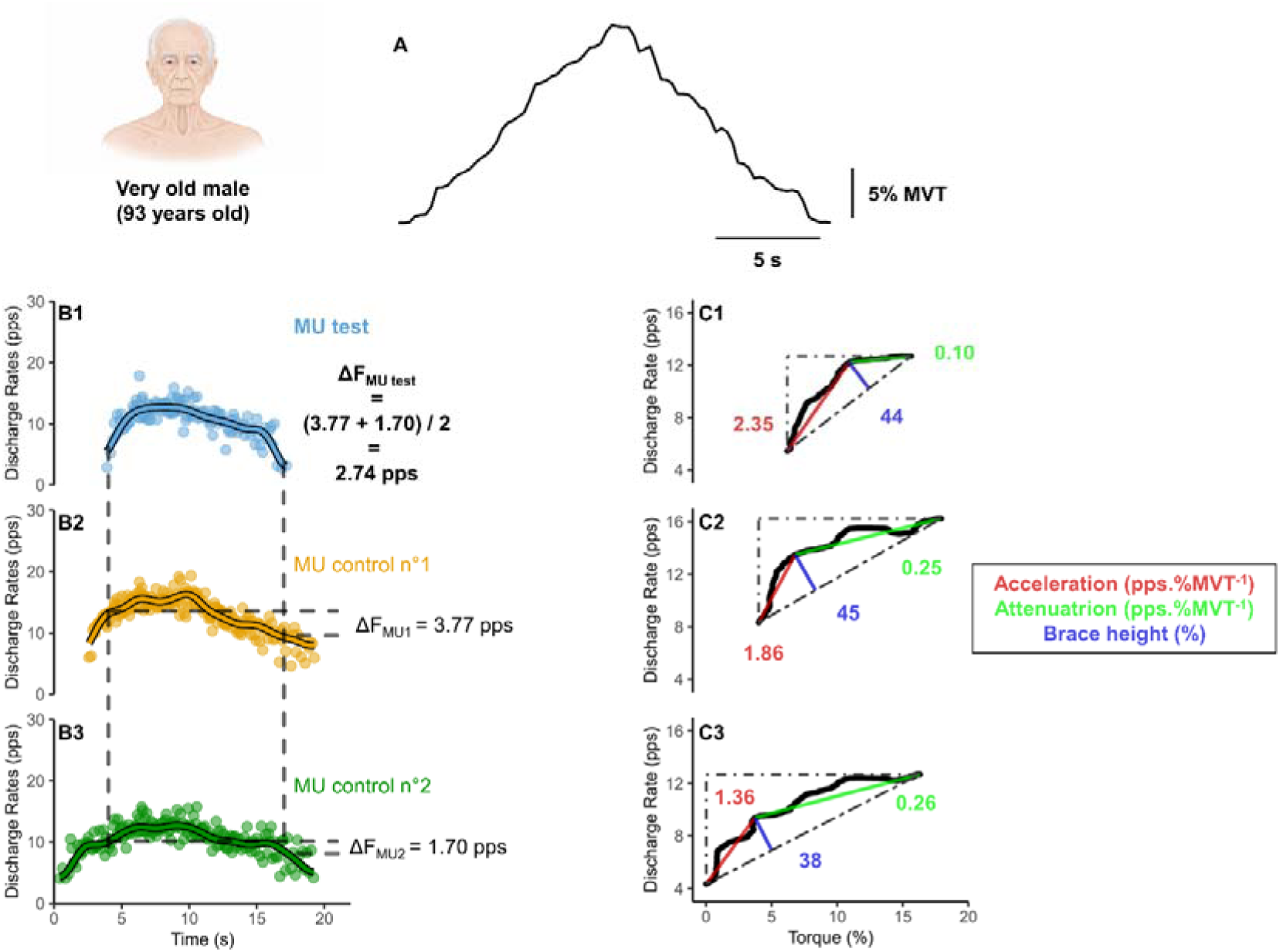
Typical trace of torque profile from one very old adult during a triangular shaped-ramp contraction at 20% of maximal voluntary force (panel A), illustration of paired MU analysis for computation of ΔF (panel B) and quasi-geometric analysis for brace height, acceleration and attenuation calculation (panel C). The left section (panels B1 to B3) shows smoothed discharge rate (solid lines outlines in black) and raw instantaneous discharge rates (dots) from three MUs identified during the triangular shaped-ramp contraction. The “test” MU is represented in blue (panel B1) and the two “control” MUs allowing to construct pairs are represented in yellow (panel B2) and green (panel B3). For each pair, the corresponding ΔF value is provided and were then averaged to obtain a single value for one test unit. The right section (panels C1 to C3) shows the discharge rate- torque relationship of the same MUs presented in left section allowing to extract brace height (in blue), acceleration slope (in red) and attenuation slope (in green). MU: Motor unit; MVT: Maximal voluntary torque; PPS: pulse per second.

#### 2.2.1) Experimental setting

Participants were seated comfortably in an isokinetic dynamometer (Biodex System 3, Biodex Medical System Inc., Shirley, USA) with the right foot securely strapped to the footplate using Velcro straps wrapped around the foot. The center of rotation of the dynamometer was aligned with the lateral malleolus. The ankle, knee and hip angles were set at 90°, 120° and 130° respectively (180° full extension). The trunk and hip were firmly strapped to the seat with belts to limit extraneous movements of the upper body. Dorsiflexion torque and EMG signals were acquired at 2048 Hz using multichannel acquisition system (EMG-Quattrocento, 400-channel EMG amplifier, OTBioelettronica, Torino, Italy).

#### 2.2.2) High-density electromyography recordings

High-density surface electromyograms were recorded using two 64-channels electrode grids (8-mm interelectrode distance; GR08MM1305, OTBioelettronica, Torino, Italy) attached to the skin with bi-adhesive foam filled by conductive paste (AC Cream, Spes Medica, Genoa, Italy). The two grids were placed in series along the muscle. The location of the *tibialis anterior* was identified through palpation before the placement of the grids. To ensure good skin-electrode contact, the grids were secured to the leg using an elastic strap wrapped around the limb. Before placing the grids, the skin was shaved and cleaned with an abrasive paste (Everi, Spes Medica, Genoa, Italy). EMG signals were acquired using OTBioLab+ software (OTBioelettronica, Torino, Italy) and were amplified (× 150) and band-pass filtered (10-500 Hz) before being stored for offline analysis. A reference electrode (30 × 24 mm; Kendall H124SG; Cardinal Health; Dublin; Ireland) was placed on the right patella while a ground electrode (strap electrode dampened with water; WS2, OTBioelettronica, Torino, Italy) was positioned around the right ankle. The *tibialis anterior* was chosen because (i) it is known to allow the identification of a large number of MUs compared to other muscles such as *soleus*, *biceps* or *triceps brachii* (Hassan et al. 2021; Orssatto et al. 2021, 2022b), which is particularly relevant for females, for whom the identification of MUs is more challenging than for males (Del Vecchio et al. 2020; Taylor et al. 2022; Jenz et al. 2023); (ii) its important involvement in postural stability and locomotion (Hernández-Guillén et al. 2021; Perera et al. 2021) and (iii) its sensitivity to detect changes in spinal motor neuron excitability in older adults (Orssatto et al. 2021).

### 2.3) Functional assessments

Old and very old participants performed a 6-minute-walk-test (6MWT) and a Timed-Up-and-Go (TUG) test to assess functional capacities. The 6MWT consisted of covering the longest distance possible in 6 min in a corridor of 30 m (Crapo et al. 2002). Standardized encouragements were given every minute. During the TUG test, participants were asked to stand up from a standard chair, walk a distance of 3 m at a conformable pace, turn around, walk back and sit down (Podsiadlo and Richardson 1991). The timer started when the participants moved their back form the chair and was stopped when the participants were seated with their back on the back of the chair.

### 2.4) Accelerometry

An accelerometer (GT3X-BT; ActiGraph, Pensacola, USA) was given to each participant to objectively assess physical activity. The accelerometer was worn on the non-dominant wrist for 7 consecutive days in free-living condition during waking hours and was removed during nighttime. Acceleration data were recorded at 60 Hz and analyzed using 60-sec epochs. Wear time was validated when a participant wore the accelerometer during at least 10 h/day for a minimum of 4 days including 2 week days and 2 weekend days. Participants who did not meet these criteria were excluded from accelerometry-based analysis (1 young adult, 2 old adults and 2 very old adults). Accelerometer data were processed using ActiLife software (version 6.13.4, ActiGraph, Pensacola, USA) and average daily step count (steps.day^-1^) was extracted for subsequent analysis.

### 2.5) Data analysis

#### 2.5.1) Decomposition of electromyographic signals

The open-source software MUEdit (Avrillon et al. 2024) was used to decompose EMG signals. MUEdit performs EMG decomposition using a blind-source separation approach [i.e., fast Independent Component Analysis (Hyvärinen 1999)] allowing the extraction of individual MU pulse trains from which discharge times were identified. After decomposition, MU pulse trains were screened for quality. Low-quality MUs were excluded and the remaining ones were manually edited when needed (Del Vecchio et al. 2020; Hug et al. 2021). Then, duplicate MUs within- and between grids were identified and removed. MUs could not be identified in 1 young female. Two old females, 1 old male and 4 very old females were excluded *a posteriori* (i.e., during data analysis) because excessive force fluctuations compromised the quality of the triangular-shaped ramp contractions. Therefore, results have been obtained from 29 young (14 women), 26 old (13 women) and 24 very old participants (10 women).

#### 2.5.2) Motor unit discharge characteristics and motor unit action potentials

Instantaneous discharge rates were smoothed using support vector regression (Beauchamp et al. 2022) with additional Matlab scripts and functions. MU peak discharge rate was considered as the maximal value obtained from the smoothed discharge rate. MU recruitment and derecruitment thresholds were computed as the relative torque level (i.e., % MVC) at the time when the MU started and stopped to discharge action potentials, respectively. MU discharge rate at recruitment and derecruitment were defined as the first and last values of the smoothed discharge rate, respectively (Škarabot et al. 2025). MU discharge rate modulation was measured as the difference between the peak discharge rate and the discharge rate at recruitment. MU action potentials (MUAPs) waveforms were reconstructed using spike-triggered averaging of bipolar EMG signals based on MU discharge timings (Farina et al. 2002). For each MU, EMG segments centered on each spike (± 25 ms) were averaged. MUAP amplitude was quantified as the mean peak-to-peak value of the five channels exhibiting the largest MUAP amplitudes.

#### 2.5.3) Paired motor unit technique

We used the paired MU technique to estimate the PICs prolongation effect (Gorassini et al. 1998, 2002). This approach quantifies recruitment-derecruitment hysteresis by pairing a lower-threshold MU (namely the control unit) with a higher-threshold MU (namely the test unit) (Figure 1B). In this context, the smoothed discharge rate of the control unit is used as a proxy for changes in net synaptic input to the motor neuron pool. For each MU pair, the PIC-related hysteresis was quantified by calculating the difference in the discharge rate of the control unit at the time of recruitment and derecruitment of the test unit. This difference, named as ΔF, is considered proportional to the prolongation effect of PICs on motor neuron firing (Gorassini et al. 2002). Only MU pairs meeting the following established criteria were retained for analysis (Hassan et al. 2020) : (i) the test MU had to discharge for a minimum duration of 2 s; (ii) the test MU were required to be recruited at least 1 s after the control MU to ensure full activation of PICs; (iii) the test MU were required to be derecruited at least 1.5 s before the control MU to avoid overestimation of ΔF; (iv) the smoothed discharge rate profiles of the test and control MUs were required to be strongly correlated (i.e., coefficient of determination r^2^ ≥ 0.7) to ensure a large proportion of common synaptic input and (v) the control MU had to modulated its discharge rate by ≥ 0.5 pps during the active period of the test MU to ensure that its discharge rate remained sensitive to changes in synaptic input (i.e., remained a valid proxy for net synaptic input). ΔF values were computed for each eligible control-test MU pair and then averaged to obtain one representative ΔF value per test MU. Test MUs forming fewer than two control-test pairs were excluded to reduce the influence of single-pair variability ensuring a robust estimate of ΔF.

Additionally, we computed normalized ΔF value which is suggested to complete traditional ΔF value when control unit discharge rates differ between populations (Škarabot et al. 2025). For each eligible pair, ΔF was normalized to the maximal discharge rate modulation of the control MU, defined as the difference between discharge rate of the control MU at test MU recruitment and the discharge rate at control MU derecruitment.

#### 2.5.4) Quasi-geometric approach

To estimate neuromodulatory and inhibitory contributions to MU discharge at the single MU level, we applied a quasi-geometric approach quantifying non-linearities in the relationship between MU discharge rate and torque output during the ascending phase of the ramp contractions (Beauchamp et al. 2023). This method characterizes the deviation of MU discharge behavior from a theorical linear pattern from recruitment to peak discharge rate (Figure 1C). For each MU, the smoothed discharge rate was plotted as a function of the corresponding torque produced during the ramp-up phase of the contraction. A theoretical linear trajectory was then defined by fitting a straight line between the discharge rate at MU recruitment and the peak discharge rate achieved during the contraction (Figure 1C). The maximum orthogonal distance between this linear fit and the smoothed discharge rate trajectory was quantified and defined as brace height. To account for differences in discharge rate range across MUs and to ensure that brace height reflected relative deviations from linearity, brace height was normalized to the height of a right triangle whose hypotenuse corresponded to the linear segment connecting recruitment and peak discharge rate. Higher normalized brace height values are interpreted as reflecting stronger neuromodulatory input (Beauchamp et al. 2023; Chardon et al. 2024). In addition to brace height, the quasi-geometric approach allows segmentation of the MU discharge trajectory into two distinct temporal phases : (i) the acceleration phase extended from MU recruitment to the point at which brace height occurred and reflects the initial rapid increase in discharge rate following recruitment, which is thought to reflect PICs-related amplification of synaptic input from the early activation of PICs and (ii) the attenuation phase extended from the brace height point to peak discharge rate and corresponds to a region where further increases in torque result in a reduced rate of increase in discharge, likely reflecting PICs saturation and the influence of inhibitory mechanisms. For each MU, the slope of the discharge rate trajectory was calculated separately for the acceleration and attenuation phases. Several exclusion criteria were applied to ensure reliable estimation of the quasi-geometric metrics: (i) negative slope during acceleration phase, (ii) negative brace height prior to normalization, and (iii) peak discharge occurring after the peak of torque profile. In addition, because an incomplete discharge trajectory may artificially truncate the attenuation phase and bias brace height estimation, MUs with attenuation phase shorter than 2 s were excluded (Goreau et al. 2025; Bontemps et al. 2026).

### 2.6) Statistical analyses

For all variables, normality of residuals was assessed using Shapiro-Wilk test, and homogeneity of variance using Levene’s test. MVT, MVT normalized to body mass, physical activity (i.e., steps.day^-1^) and 6MWT performance were analyzed using linear models (lm package) including age, sex and age × sex interaction as fixed effect. For TUG test performance, which violated the normality assumption, a nonparametric analysis based on the Aligned Rank Transform [ARTool package (Wobbrock et al. 2011)] was applied. Post-hoc comparisons were performed using Tukey’s HSD test when appropriate.

To assess changes in MU discharge characteristics (i.e., peak discharge rate, recruitment and derecruitment thresholds, discharge rate at recruitment and derecruitment, discharge rate modulation, MUAP amplitude, ΔF and normalized ΔF, brace height, acceleration slope, and attenuation slope) across age and sex, linear mixed-effects models were performed. Prior to analyses, outliers (> 3 interquartile range) were removed at the MU level for each metric. Linear mixed-effects models were applied to individual MU values, with age (young, old, and very old), sex (males, females) and age × sex interaction as fixed effects and a random intercept for each participant: Variable ∼ age × sex + (1|participant) using lmer package (Kuznetsova et al. 2017).

When significant effects were detected using the Satterthwaite’s approximation for degrees of freedom, Tukey post hoc correction was applied for pairwise comparisons. Estimated marginal means (EMM) and 95% confidence intervals (CI 95%) were computed using the emmeans package. Data are presented as EMM and CI 95% within the table and figures. All statistical procedures and figure generation were performed on RStudio (version 2025.09.2) (R Core Team 2025). Statistical significance was at p<0.05. Database and R code can be found at: https://osf.io/3maut/

## **III)** RESULTS

### 3.1) Maximal voluntary torque, physical activity and functional capacities

Participants’ characteristics are summarized in Table 1. Females showed lower absolute MVT than males (p<0.001) without main effect of age (p=0.210) or age-by-sex interaction (p=0.082). When normalized to body mass, MVT was also lower in females (p=0.002) and was influenced by age (p=0.005) without age-by-sex interaction (p=0.099). Specifically, normalized MVT was significantly lower in old (p=0.030) and very old (p=0.006) compared to young adults, without significant difference between old and very old adults (p=0.802). Physical activity level (expressed as steps.day^-1^) was significantly lower in very old compared to old adults (p<0.001), without significant difference between young and both old (p=0.216) and very old adults (p=0.061). Old adults performed better at the 6MWT (p<0.001) and the TUG test (p<0.001) than very old adults.

**Table 1.** Participants’ characteristics. BMI: Body mass index; MMSE: Mini-mental state examination; MVT: Maximal voluntary torque; TUG: Timed-up and go; 6MWT: 6 minutes walking test. Data are presented as estimated marginal mean and 95% CI lower and upper limits. Significantly different from young adults: ** p<0.01; *** p<0.001. Significantly different from old adults: ### p<0.001. Significantly different from males: $ p<0.05; $$ p<0.01; $$$ p<0.001.

| Characteristics | Young males<br>(n=15) | Young females<br>(n=14) | Old males<br>(n=13) | Old females<br>(n=13) | Very old males<br>(n=14) | Very old females<br>(n=10) |
| --- | --- | --- | --- | --- | --- | --- |
| **Anthropometric data** |  |  |  |  |  |  |
| Age (years) | 26.2 [24.0-28.4] | 25.6 [23.4-27.9] | 70.5 [68.1-72.8] *** | 71.2 [68.9-73.6] *** | 84.3 [82.0-86.6] ***-### | 84.1 [81.4-86.8] ***-### |
| Height (cm) | 173.9 [170.7-177.2] | 166.1 [162.8-169.5] \$\$\$ | 171.8 [168.4-175.3] | 161.8 [158.4-165.3] \$\$\$ | 170.6 [167.2-173.9] | 161.2 [157.3-165.1] \$\$\$ |
| Weight (kg) | 67.9 [63.1-72.7] | 58.4 [53.4-63.3] \$\$\$ | 73.8 [68.5-78.9] | 61.8 [56.6-66.9] \$\$\$ | 73.5 [68.6-78.5] | 60.5 [54.6-66.4] \$\$\$ |
| BMI (kg.m<sup>-2</sup>) | 22.4 [21.0-23.8] | 21.2 [19.7-22.6] \$ | 24.9 [23.3-26.4] ** | 23.5 [22.0-25.1] **-\$ | 25.2 [23.8-26.7] ** | 23.3 [21.5-25.1] **-\$ |
| **Questionnaires, physical activity and functional capacities** |  |  |  |  |  |  |
| MMSE (0-30) | - | - | 28.6 [28.0-29.3] | 28.5 [27.8-29.2] | 28.2 [27.5-28.9] | 28.4 [27.6-29.1] |
| Steps/day (n) | 11082<br>[9659-12506] | 11356<br>[9933-12780] | 11923<br>[10385-13460] | 13027<br>[11490-14565] | 9171<br>[7693-10648] ### | 9775<br>[8091-11459] ### |
| TUG test (s) | - | - | 7.1 [6.0-8.1] | 7.2 [6.2-8.3] | 9.7 [8.7-10.7] ### | 9.9 [8.7-11.1] ### |
| 6MWT (m) | - | - | 601.8 [561.4-642.2] | 547.1 [506.7-587.5] | 473.7 [434.8-512.7] ### | 457.1 [411.0-503.2] ### |
| MVT (Nm) | 40.8 [37.2-44.3] | 32.8 [29.1-36.5] \$\$\$ | 42.7 [38.9-46.5] | 27.0 [23.2-30.8] \$\$\$ | 37.1 [33.4-40.7] | 28.7 [24.3-33.1] \$\$\$ |
| MVT (Nm.kg<sup>-1</sup>) | 0.60 [0.55-0.65] | 0.56 [0.51-0.62] \$\$ | 0.58 [0.53-0.64] *** | 0.44 [0.39-0.50] ***-\$\$ | 0.51 [0.46-0.57] *** | 0.48 [0.42-0.54] ***-\$\$ |

### 3.2) Motor unit identification and discharge characteristics

A total of 2325 MUs (22.9 ± 13.5 per participant in young adults, 33.2 ± 15.0 per participant in old adults and 33.3 ± 11.1 per participant in very old adults) were identified during the triangular shaped- ramp contractions. Supplementary material S1 details the number of detected MUs, number of MU pairs, number of test MUs used for ΔF and normalized ΔF calculation as well as number of MU retained for geometric metrics computation across age and sex groups.

Peak discharge rate was higher in young adults compared to both old (p=0.003) and very old adults (p=0.002), whereas no differences were observed between old and very old adults (p=0.989), independent of sex (p=0.115) (Figure 2A). Similarly, discharge rate at recruitment was higher in young compared to both old and very adults (both p<0.001) without difference between old and very old adults (p=0.906), independent of sex (p=0.464) (Figure 2D). There was a significant age-by-sex interaction on derecruitment threshold (p=0.045). Specifically, young females presented higher values compared to old (p=0.019) and very old females (p=0.002), while no differences were observed in males (Figure 2C). MUAP amplitude was higher in old (p=0.010) and very old adults (p=0.018) compared to young adults, whereas no differences were observed between old and very old adults (p=0.991), independent of sex (p=0.085) (Figure 2G). Finally, no effects of age and sex were reported for recruitment threshold (Figure 2B), discharge rate at derecruitment (Figure 1E) and discharge rate modulation (Figure 2F).

**Figure 2.**
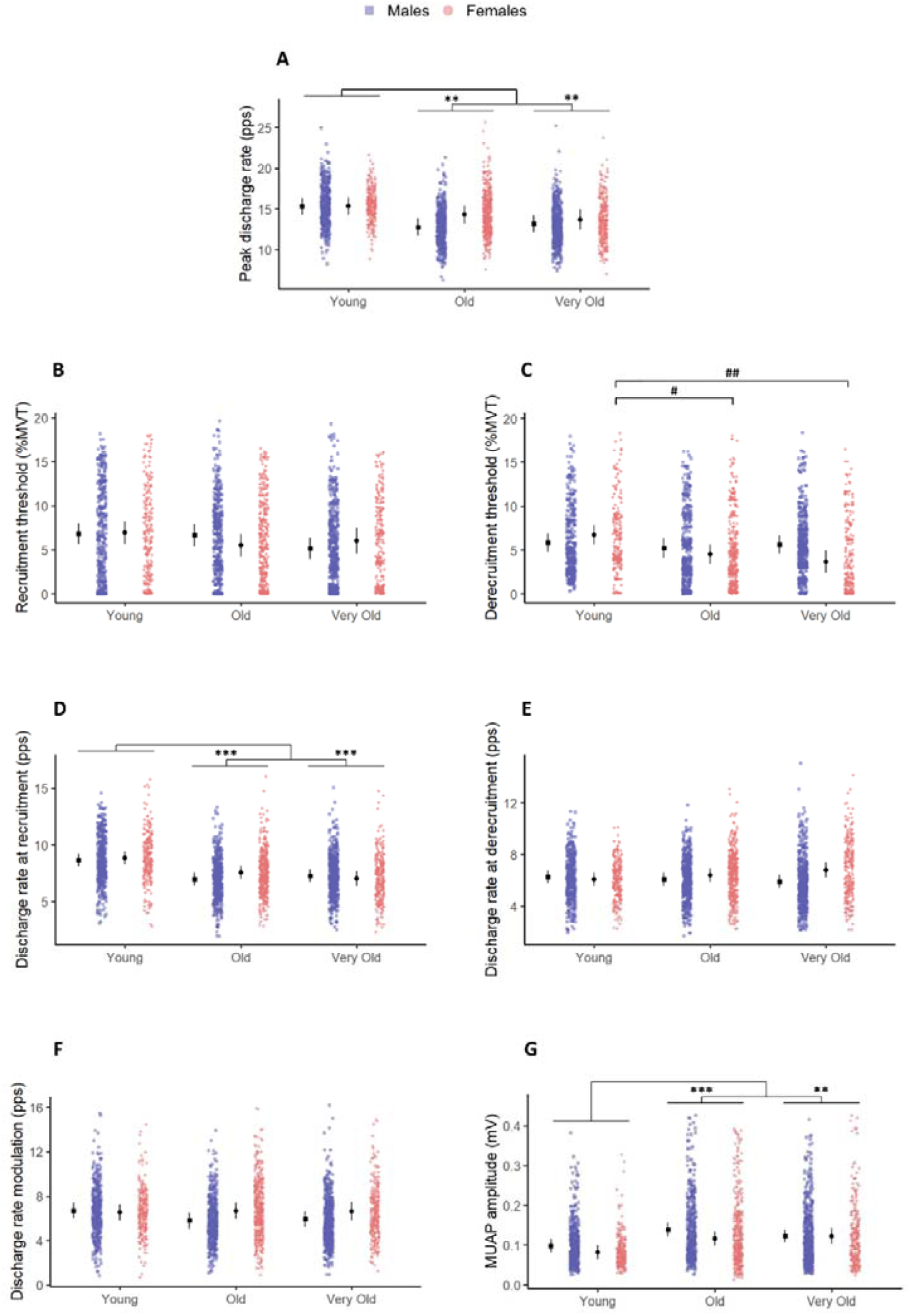
Differences between age groups and between sex for peak discharge rate (panel A), recruitment threshold of the identified MUs (panel B), derecruitment threshold of the identified MUs (panel C), discharge rate at recruitment (panel D) and discharge rate at derecruitment (panel E), discharge rate modulation (i.e., peak discharge rate – discharge rate at recruitment) (panel F), MUAP amplitude (panel G). Black circles and lines represent estimated marginal means and 95% confidence intervals, respectively. Symbols in color (blue for males and pink for females) represent individual MU values. MVT: Maximal voluntary torque; MUAP: Motor unit action potential; PPS: pulse per second. Significantly different from young: ** p<0.01; *** p<0.001. Significant difference between age groups for females: # p<0.05; ## p<0.01.

### 3.3) Estimates of PICs prolongation effect

For ΔF and normalized ΔF, six participants were excluded from the analyses (1 young female, 1 old female, 1 old male, 1 very old female and 2 very old males) because it was not possible to achieve all the assumptions required in the paired MU analysis. Therefore, results were obtained from 28 young adults (13 females), 24 old adults (12 females) and 21 very old adults (9 females).

ΔF was higher in young adults compared to both old and very old adults (all p<0.001), whereas no differences were observed between old and very old adults (p=0.811) (Figure 3A). A significant age-by-sex interaction was reported for normalized ΔF (p<0.001) (Figure 3B). Specifically, young males showed higher normalized ΔF values than both old (p<0.001) and very old males (p=0.019) without significant difference between old and very old males (p=0.268). In females, lower values were observed in very old females compared to both young and old females (both p<0.001), while no difference was reported between young and old females (p=0.247).

**Figure 3.**
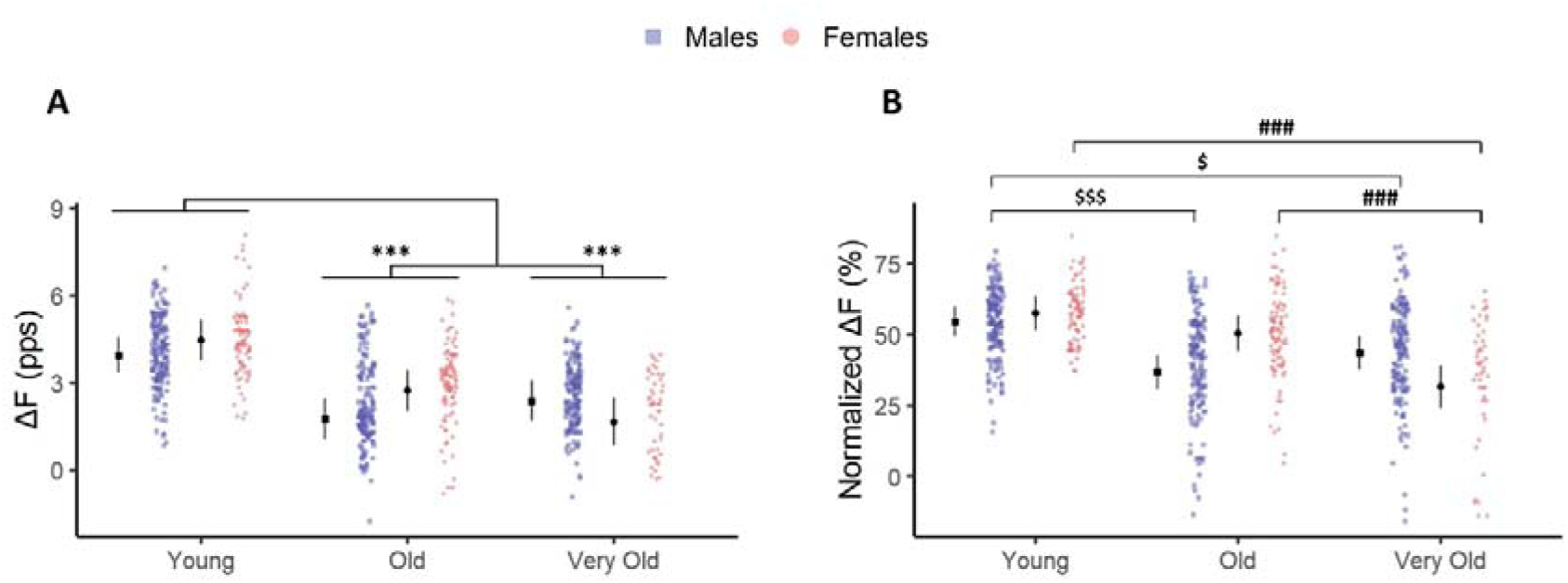
Differences between age groups and between sex for ΔF (panel a) and normalized ΔF (panel b). Black circles and lines represent estimated marginal means and 95% confidence intervals, respectively. Symbols in color (blue for males and pink for females) represent individual test MU values. PPS: pulse per second. Significantly different from young: *** p<0.001. Significant difference between age groups for males: $ p<0.05; $$$ p<0.001. Significant difference between age groups for females: ### p<0.001.

### 3.4) Quasi-geometric approach

For quasi-geometric analyses, two participants (1 young male and 1 very old females) were excluded from the analyses because it was not possible to ensure the assumptions required for at least one MU, i.e., positive slope during acceleration phase, positive brace height and peak discharge occurring before the peak of torque profile. Therefore, results were obtained from 28 young (14 females), 26 old (13 females) and 23 very old adults (9 females).

No effect of age, sex or age-by-sex were reported for brace height (Figure 4A), acceleration slope (Figure 4B) and attenuation slope (Figure 4C).

**Figure 4.**
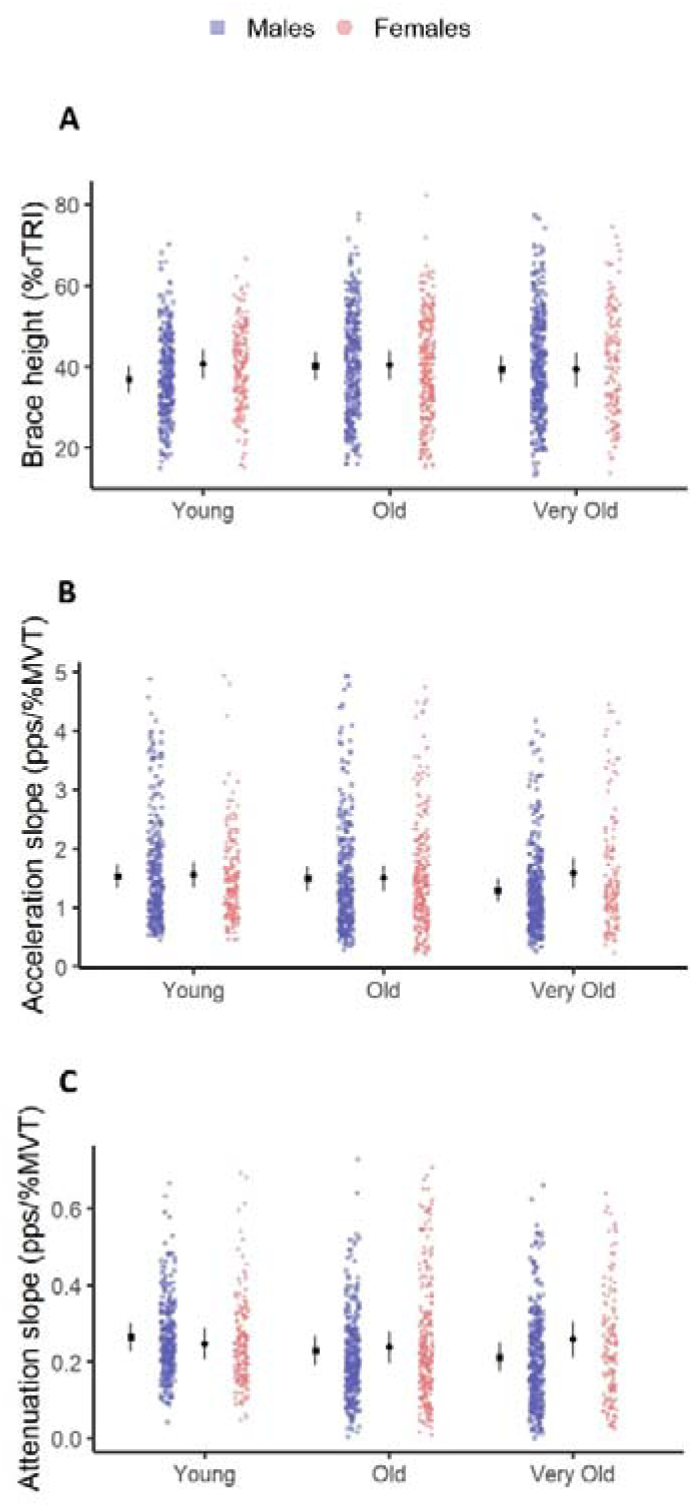
Differences between age groups and between sex for brace height (panel A), acceleration slope (panel B) and attenuation slope (panel C). Black circles and lines represent estimated marginal means and 95% confidence intervals, respectively. Symbols in color (blue for males and pink for females) represent individual MU values. MVT: Maximal voluntary torque; PPS: pulse per second; %rTRI: percentage of right triangle.

## **IV)** DISCUSSION

The present study provides new insight into how spinal motor neuron function evolves with advanced aging. Our findings indicate that major age-related changes in MU discharge behavior and PICs-related estimates are already established by the seventh decade and do not appear to worsen further in healthy active adults over 80 years of age. Importantly, this apparent stabilization at very old age was accompanied by evidence of distinct age-related trajectories between males and females when PICs-related hysteresis was expressed relative to the available discharge rate modulation. Finally, the preservation of indices of neuromodulatory and inhibitory influences indicate that the lower PICs-related hysteresis observed with aging is unlikely to result from a reduced monoaminergic drive. Together, these findings suggest that age-related changes in spinal motor neuron excitability do not follow a continuous decline and highlight both the nonlinearity and the potential sex dependence of neuromuscular aging.

### 4.1) Age-related changes in rate coding

Both peak discharge rate and discharge rate at recruitment were lower in old and very old adults compared to young adults, consistent with previous findings (for review, see Orssatto et al. 2022a). The present results extend them by showing that this age-related difference does not further increase beyond 80 years of age, at least for the tibialis anterior muscle. Despite the lower peak discharge rate in old and very old adults compared to young adults, absolute MVT was similar across age groups, regardless of sex. This may be explained by the slower contractile properties with aging (Doherty and Brown 1997), which enhance temporal summation and thereby lower discharge rate required to produce a given force (Botterman et al. 1986; Fuglevand et al. 1999). A complementary explanation lies in the denervation-reinnervation cycle of muscle fibers that occurs over the lifespan resulting in fewer but larger MU (Gordon et al. 2004). In the present study, this interpretation is supported by the greater MUAP amplitude observed in old and very old adults compared to young individuals, as larger MUs are generally characterized by greater action potential amplitudes (Hakansson 1956; Pope et al. 2016). Such remodeling may partially compensate for reduced discharge rates by increasing the force contribution of individual MUs, thereby preserving MVT. Nonetheless, the lower MVT normalized to body mass in old and very old adults suggests that this compensation remains limited.

### 4.2) Potential mechanisms underlying age-related changes in motor unit behavior

Age-related changes in MU behavior may partly arise from alterations in the intrinsic excitability of spinal motor neurons. We observed significantly lower ΔF values in older adults, in agreement with previous studies (Hassan et al. 2021; Orssatto et al. 2021, 2022b; Guo et al. 2024). The lower prolongation effect of PICs in older adults has been suggested to result, at least in part, from a reduced monoamine release (i.e., serotonin and noradrenaline) at synapses on spinal motor neurons and/or a decreased receptors sensitivity (Orssatto et al. 2023). The absence of group differences in brace height, a metric thought to primarily reflect neuromodulation (Beauchamp et al. 2023; Chardon et al. 2024), does not support a major impairment in monoaminergic drive with aging, even in very old adults. In addition, the lower ΔF values observed in old and very old adults, together with an acceleration slope unaffected by age, might suggest that aging preferentially affects mechanisms related to the prolongation rather than the amplification effect of PICs. Such a dissociation could potentially involve the distinct voltage-sensitive channels generating PICs, which may contribute differently to motor neuron discharge behavior (Heckman et al. 2005; Binder et al. 2020). Specifically, previous studies have suggested that persistent sodium currents contribute primarily to spike initiation and rapid discharge acceleration, whereas persistent calcium currents exhibit slower and more sustained activation properties and are considered major contributors to discharge hysteresis and self-sustained firing (Lee and Heckman 1998; Li et al. 2004). Therefore, our results suggest that aging is not associated with a major alteration of PICs amplification or neuromodulatory drive, at least in active older adults.

At first glance, the concomitant absence of differences in attenuation slope also suggests that inhibitory patterns may be unchanged with aging. However, attenuation slope is derived exclusively from discharge non-linearities during the ascending phase of the ramp contraction, whereas ΔF reflects the hysteresis of MU discharge across both ascending and descending phases. This distinction raises the possibility that age-related changes in inhibitory input may occur preferentially during the descending phase of the contraction in old and very old adults. Several MU discharge characteristics support this interpretation: (i) preserved discharge rate modulation across groups, indicating a maintained ability to increase discharge rate during the ascending phase in old and very old individuals, and (ii) higher discharge rate at recruitment in young adults, while discharge rate at derecruitment was similar across groups suggesting distinct regulatory mechanisms between the ascending and descending phase during ramp contractions. Such phase-dependent inhibitory modulation could explain the preserved attenuation slope despite lower ΔF values with aging.

Additional indirect evidence for a greater role of inhibitory mechanisms comes from the higher levels of antagonist coactivation typically observed in older adults (Macaluso et al. 2002; Tracy and Enoka 2002), since co-contraction has been shown to reduce ΔF through increased reciprocal inhibition (Gomes et al. 2024). Interestingly, Gomes et al. (2024) reported a reduction in ΔF without changes in attenuation slope, in line with the present findings. Such a dissociation, whereby ΔF is reduced without detectable changes in quasi-geometric metrics has also been reported in response to experimental pain and in sarcopenic individuals (Hug et al. 2025; Orssatto et al. 2025). Therefore, the absence of age-related difference in attenuation slope in the present study should be interpreted with caution rather than as evidence of preserved inhibitory pattern. Recently, Connelly et al. (2026) reported a lower attenuation slope in older adults with a preserved brace height, suggesting an altered pattern of excitation-inhibition coupling rather than a reduced neuromodulatory drive.

### 4.3) Absence of further decline in very old adults

Contrary to our main hypothesis, the lower discharge rate and ΔF observed in old adults were not further exacerbated in very old adults, suggesting that these changes may plateau earlier in the aging process, at least in active and healthy older adults. Overall, it is likely that a substantial part of the impairments already occurs by the seventh decade, leading to an early decrease in ΔF and explaining why further chronological aging may not be associated with a lower estimated contribution of PICs to MU discharge behavior. Moreover, the lower ΔF observed in old and very old adults compared to young adults, together with the absence of any difference in quasi-geometric metrics between the two older groups, suggest that the balance between neuromodulatory and inhibitory contributions to ΔF does not shift from old to very old age.

Several factors may also account for this apparent stabilization. First, because all parameters were estimated during ramp contractions at 20% MVT, the MUs analyzed in the present study predominantly exhibited a low recruitment threshold. Such MUs are recruited during most activities of daily living and may therefore be relatively spared by the aging process. Consistently, previous studies showed that age-related differences were more pronounced at higher contraction intensities for both motor unit discharge rate (Orssatto et al. 2022a) and ΔF (Connelly et al. 2026). Our estimates were thus obtained under conditions in which age-related differences in MU behavior are the least expressed, which may have limited our ability to detect additional differences between old and very old adults.

Second, the very old adults involved in our study likely represent a relatively high-functioning subset of this age group, potentially exhibiting attenuated neuromuscular impairments compared to the general population of this age. Importantly, Orssatto et al. (2025) reported that ΔF in the tibialis anterior was substantially greater in both non-sarcopenic older adults and master athletes compared to sarcopenic older adults, suggesting that intrinsic motor neuron excitability may not be further modulated among very old adults with preserved neuromuscular function. When compared to normative reference values, very old adults of the present study showed (i) an important distance covered during the 6MWT (465 m; Otadi and Malmir 2026); (ii) a short time to perform the TUG test (9.8 s; Long et al. 2020) and (iii) a high level of physical activity (9472 steps.day^-1^; Tudor-Locke et al. 2013). As such, our sample may not fully reflect the broader very old population, particularly individuals with greater functional impairments or those living in institutionalized settings. Future studies should aim to include more representative cohorts, despite the additional methodological challenges this may entail, like medication use or reduced ability to perform triangular-shaped ramp contraction.

### 4.4) Distinct age-related trajectories of PICs in males and females

Regardless of age group, we did not observe any effect of biological sex on ΔF values or geometric-derived metrics. This contrasts with the few studies that have explored the influence of biological sex on PICs, which have been restricted to young populations (i.e., adolescents and young adults), and reported greater ΔF values in females compared to males, suggesting higher contribution of PICs in females (Jenz et al. 2023; Yacyshyn et al. 2025). One possible explanation for the absence of sex-related differences reported in the current study may lie in the contraction intensity used to estimate ΔF during isometric triangular-shaped ramp contraction (i.e., 20% MVT). For instance, findings from Jenz et al. (2023) suggesting that estimates of PICs were larger in females than in males were obtained from contractions performed at 30% MVT. Importantly, Yacyshyn et al. (2025) reported that in young adults ΔF was higher in males than in females at 10% MVT, similar between both sexes at 20% MTV and became higher in females than in males at 30% MVT, suggesting a substantial influence of contraction intensity on sex-related differences on PICs prolongation effect. These findings suggest that differences between males and females may become more apparent at higher levels of voluntary drive and that the use of a moderate contraction intensity in the present study may have limited our ability to detect potential effect of biological sex on ΔF values.

Our results obtained on normalized ΔF, which controls for differences in control unit discharge rate between populations, suggest distinct age-related trajectories between males and females. Indeed, in males, normalized ΔF values were already lower in old than in young adults, with no further difference in very old adults. Conversely, in females normalized ΔF did not differ significantly between young and old adults, but was lower only at very old age. These findings suggest that the age-related decline in PICs prolongation effect may occur earlier in males compared to females. In the literature, it is noteworthy that most studies reporting lower PICs estimates in older adults have predominantly included male participants (Hassan et al. 2021; Guo et al. 2024). Our findings therefore raise the possibility that the inclusion of females may attenuate the magnitude of differences typically observed between young and old adults and outline the importance to include females to better understand the effect of aging on PICs amplitude. Future studies are required to clarify the mechanisms underlying these distinct trajectories between males and females with aging.

### 4.5) Future directions

The results of the present study were obtained from the tibialis anterior muscle, selected for its high MU yield, at a single contraction intensity. Although this muscle is sensitive to detect changes in spinal motor neurons excitability in older adults, its size and strength appear relatively well preserved with aging (Simoneau et al. 2005; Naruse et al. 2023; Fuchs et al. 2023). Extending these measurements to muscles exhibiting greater age-related differences in muscle mass and strength, and to higher contraction intensities, would therefore clarify how far the present findings can be generalized. Moreover, whether the absence of further decline beyond the seventh decade also applies to inactive or deconditioned very old adults remains to be determined. Finally, the cross-sectional design, based on comparisons between independent age groups, may mask some age-related changes in PICs due to interindividual variability. Longitudinal studies are needed in the future to better characterize individual changes in PICs with aging.

## **V)** CONCLUSION

Our results suggest that spinal motor neuron excitability is altered with aging, but that these alterations do not follow a continuous decline up to very old age and may differ between males and females. These age-related alterations do not appear to result from a lower neuromodulatory drive, and might rather involve changes in the inhibitory control of motor neuron activity. Overall, these findings highlight the non-linear and potentially sex-dependent nature of spinal motor aging.

## Abbreviations

ANOVA: Analysis of variance BMI: Body mass index
CI: Confidence interval
EMM: Estimated marginal mean
HDsEMG: High-density surface electromyography
MMSE: Mini-mental state examination
MVT: Maximal voluntary torque
MUAP: Motor unit action potential
MU: Motor unit
PICs: Persistent inward currents
PPS: Pulse per second
SE: Standard error
TUG test: Timed up and go test
6MWT: Six-minute walking test
ΔF: Delta frequency

## CRediT authorship contribution statement

**Chatain C.**: Conceptualization; Data curation; Formal analysis; Investigation; Methodology; Software; Visualization; Writing – original draft; Writing – review & editing. **Hamard R.**: Writing – review & editing. **Cattagni C.**: Conceptualization; Validation; Writing – review & editing. **Roty R.**: Writing – review & editing. **Lepers R**: Writing – review & editing. **Rozand V.**: Conceptualization; Funding acquisition; Methodology; Supervision; Writing – original draft; Writing – review & editing.

## Data availability statement

Edited motor unit firing files, R code and database can be found at https://osf.io/3maut/

## Conflict of interest disclosure

The authors declare to have no financial, personal or other conflicts of interest.

## Notes

https://osf.io/3maut/

### Competing Interest Statement

The authors have declared no competing interest.

