## Supplementary material for "Estimates of persistent inward currents in human spinal motor neurons: non-linear and sex-dependent trajectories from young to very old age"

### **Supplementary material – S1**

***Supplementary table 1.*** *Number of motor units, motor unit pairs, test motor units and motor units retained for quasi-geometric approach per age group and sex. Values are presented as mean ± SD per participant, with subgroup total in parentheses. Sample size indicates the number of participants contributing to each variable. MU: Motor unit.*

|  | **Young** | | **Old** | | **Very old** | |
| --- | --- | --- | --- | --- | --- | --- |
|  | Males | Females | Males | Females | Males | Females |
| **All MUs** | **28 ± 13**  **(416)** | **18 ± 12**  **(247)** | **34 ± 13**  **(446)** | **32 ± 18**  **(416)** | **38 ± 11**  **(532)** | **27 ± 7**  **(268)** |
| *Sample size* | *n=15* | *n=14* | *n=13* | *n=13* | *n=14* | *n=10* |
| **MUs quasi-geometric approach** | **20 ± 11**  **(287)** | **12 ± 10**  **(175)** | **24 ± 14**  **(307)** | **21 ± 14**  **(275)** | **26 ± 12**  **(357)** | **16 ± 7**  **(141)** |
| *Sample size* | *n=14* | *n=14* | *n=13* | *n=13* | *n=14* | *n=9* |
| **MU pairs** | **180 ± 195**  **(2697)** | **90 ± 119**  **(1166)** | **214 ± 238**  **(2564)** | **169 ± 198**  **(2030)** | **184 ± 146**  **(2214)** | **65 ± 60**  **(582)** |
| *Sample size* | *n=15* | *n=13* | *n=12* | *n=12* | *n=12* | *n=9* |
| **Test MUs** | **11 ± 7**  **(168)** | **7 ± 5**  **(87)** | **13 ± 11**  **(158)** | **9 ± 7**  **(111)** | **14 ± 10**  **(164)** | **6 ± 5**  **(58)** |
| *Sample size* | *n=15* | *n=13* | *n=12* | *n=12* | *n=12* | *n=9* |
